# Rapid adaptive evolution of microbial thermal performance curves

**DOI:** 10.1101/2024.04.30.590804

**Authors:** Megan H. Liu, Ze-Yi Han, Yaning Yuan, Katrina DeWitt, Daniel J. Wieczynski, Kathryn M. Yammine, Andrea Yammine, Rebecca Zufall, Adam Siepielski, Douglas Chalker, Masayuki Onishi, Fabio A. Machado, Jean P. Gibert

## Abstract

Microbial respiration is a key biotic driver of climate change. Warming boosts microbial population growth, which increases biomass and respiration. This feedback might be disrupted by adaptation in thermal performance curves (TPCs) –whose shape describes how temperature drives growth. In this study, we uncover substantial genetic variation (G) in microbial intrinsic population growth rates (*r*), demonstrate a causal link between G variation in *r* and G variation in TPC shape, and show how this variation constrains r-TPC shape evolution along specific evolutionary paths across temperatures. We also uncover Gene-by-Environment (G × E) variation in *r*, which results in specific signatures in TPC shape and predictable temperature-dependent rapid TPC evolution but also lower G, which could reduce future evolutionary potential. Overall, we show how temperature-dependent evolution in a linchpin of global ecosystem function—microbial TPC shape—is determined by a combination of heritable and non-heritable variation in intrinsic growth rates.

## INTRODUCTION

Microbes play a central role in regulating the global carbon (C) cycle that controls climate change (Falkowski *et al*. 2008). Indeed, soil microbial respiration releases ∼94Pg/yr of C into the atmosphere (Stell *et al*. 2021) while microalgae fix 30-50 Pg of C/yr globally (Falkowski 1994). Global warming is expected to alter these microbial processes (IPCC 2023), but anticipating these effects requires a deeper understanding of the biotic and abiotic factors influencing microbial respiration in a warming world (Rocca *et al*. 2022; Wieczynski *et al*. 2023).

One such factor is microbial population growth, which influences total standing biomass, and hence, total microbial respiration (Savage *et al*. 2004). The thermal performance curve of intrinsic population growth rate (*r*) describes how microbial population growth changes with temperature (‘*r*-TPCs’ henceforth, Fig 1). The shape of r-TPCs is controlled by temperature-dependent metabolism: metabolism increases with temperature, and so does *r*, until an ‘optimal’ temperature (T_opt_) is reached (Fig 1a), then *r* declines as metabolic costs increase (Fig 1a; Amarasekare & Savage 2012). While *r*-TPC shape varies across species (Jacob & Legrand 2021;), unimodal shape is the norm (Wieczynski *et al*. 2021), described by shape parameters linked to thermal ecology: maximum growth rate (r_peak_), minimum and maximum temperatures for population growth (CT_min_ and CT_max_ respectively), and rates of *r*-TPC increase/decrease with temperature (E_a_, E_d_ respectively, Fig 1b). Ultimately, differences in *r*-TPC shape across species reflect divergent evolutionary trajectories in shape parameters across species and environments (Angilletta 2009).

**Figure 1.**
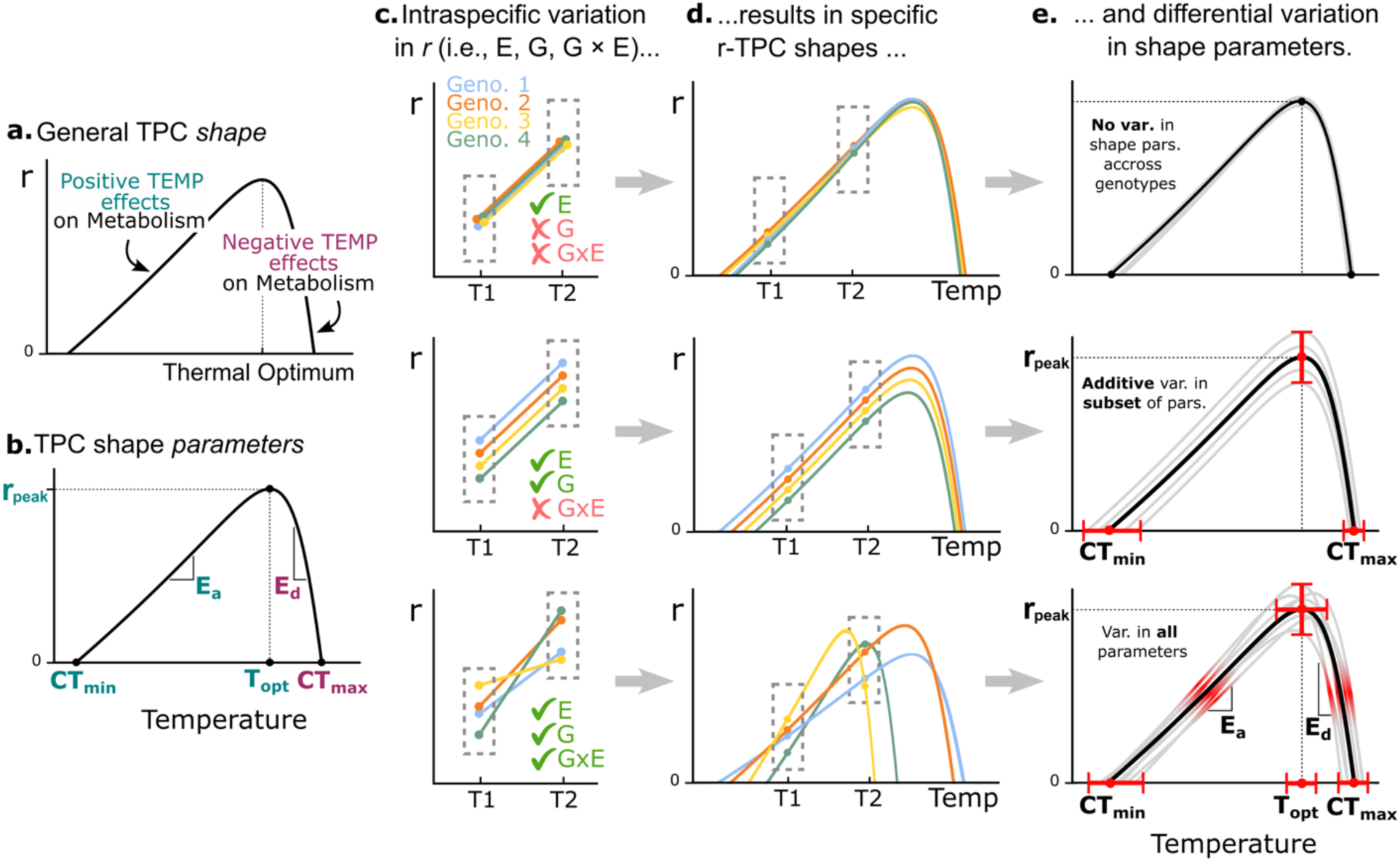
(a) General shape of the r-TPC. (b) r-TPC shape parameters. Blue-colored shape parameters, measured in this study, include r_peak_, E_a_, CT_min_ and T_opt_. (c) Top: environmental variation (E) in *r* comes from expressing different *r* across temperatures but not genotypes, and leads to overlapping reaction norms. Middle: genetic variation (G) in *r* results from expressing different *r* across genotypes, but not temperatures, and leads to additive shifts in reaction norm intercepts, but not slopes. Bottom: gene-by-environment interactions (G × E) results from patterns expression of expression of r changing across temperature, leading to shifts in slopes of reaction norms. (d) Because-rTPCs are multi-temperature reaction norms of *r*, classic quantitative genetics can explain how E, G, G × E in *r* influence r-TPC shape: E in *<u>r</u>* produces fully overlapping r-TPCs (top), G in *r* result in non-overlapping r-TPCs with genotype-specific intercepts but not slopes (middle), and G × E leads to r-TPCs that vary in intercepts and slopes (bottom). (e) These r-TPC shape signatures create variation in r-TPC shape parameters: overlapping r-TPCs show no variation in shape parameters (top), additive shifts in r-TPC intercepts lead to additive (heritable) variation in r_peak_, CT_min_, and CT_max_ (middle), while slope and intercept variation in r-TPCs creates variation in all shape parameters.

While *r*-TPC shape is expected to evolve in novel environments, anticipating this thermal adaptation is a major open question, as the r-TPC reflects a species’ ability to cope with environmental change. Thermal adaptation hinges on the evolution of intraspecific—heritable— genetic variation and selection favoring genetic variants suited to novel environments (Frankham 2005), so quantifying heritable intraspecific variation in *r*-TPC shape parameters is central to understanding *r*-TPC shape evolution (Kling *et al*. 2023). From a quantitative genetics standpoint, the *r*-TPC represents the reaction norm of the underlying measure of performance, *r*, across temperatures among genotypes (Golmulkiewicz *et al*. 2018). *r*-TPC intraspecific variation is thus inextricably linked to that of *r*. Understanding this link is key to quantifying heritable variation in *r*-TPC shape parameters and predicting their response to selection in novel environments.

To understand the link between intraspecific variation in *r* and *r*-TPC shape parameters, we developed a verbal model using classic quantitative genetics of reaction norms (Fig 1, c-e). First, environmental variation (E) in *r*, driven by differential expression of *r* across temperatures, but not genotypes, classically manifests as completely overlapping genotypic reaction norms (Fig 1C, first row). Correspondingly, genotypic *r*-TPCs should also overlap (Fig 1D, first row) as they are reaction norms extended over multiple temperatures. Overlapping *r*-TPCs have no variation in shape parameters (Fig 1e, first row), so E in *r* should not result in *r*-TPC shape parameter variation. Second, genetic variation (G) in *r*, arising from differential expression of *r* across genotypes, but not temperatures, results in “additive” shifts in the intercepts of genotypic reaction norms—but equal slopes (Fig 1C, second row). These additive shifts translate to parallel *r*-TPCs (Fig 1D, second row), with additive variation in their x-axis intercepts (CT_min_, CT_max_) and their maximum height (r_peak_), but no variation in T_opt_, E_a_, E_d_ (Fig 1E, second row). Last, gene-by-environment (G × E) interactions in *r*, which manifest as changes in intercept and slope of reaction norms across genotypes and temperatures (Fig 1C, third row), result in non-parallel *r*-TPCs (Fig 1D, third row), with greater variation in all shape parameters (Fig 1E, third row). This conceptual model therefore predicts covariation between heritable genetic variation in *r* and variation in CT_min_, CT_max_, and r_peak_, so *r*-TPCs’ CT_min_, CT_max_, and r_peak_ are more likely to be heritable and respond to selection than T_opt_, E_a_, or E_d_, which in turn suggest constraints operating on possible *r*-TPC evolutionary trajectories.

To address how *r*-TPC shape might evolve across temperatures we thus quantify E, G, and G × E variation in *r*, and test predictions from our conceptual model. In doing so we reveal that heritable variation in *r* results in substantial heritable variation in predicted shape parameters. We build upon this understanding to quantify the structure of genetic variation and covariation in shape parameters, and use this information to infer selection and possible *r*-TPC evolutionary trajectories of shape parameters across temperatures. Last, we uncover a mechanism of rapid *r*-TPC evolution in the presence of G × E variation in *r* that leads to temperature-dependent selection on *r*-TPC shape, but can erase heritable variation, possibly creating an evolutionary trap that constrains future microbial *r*-TPC evolution.

## METHODS

### Study system and genotypes

We used *Tetrahymena thermophila*, a freshwater ciliate protist found across the Northeastern United States (Zufall *et al*. 2013) and part of a cosmopolitan genus (Lynn & Doerder 2012). Protists, unicellular Eukaryotes that dominate oceanic biomass and rank third in terrestrial biomass, comprise twice the biomass of the Animal Kingdom (Bar-On *et al*. 2018) and underpin global ecosystem functioning (Gao *et al*. 2019). So, while no single species is representative of the entire group, understanding thermal adaptation in such an ecologically pivotal group is meaningful.

To do so, we sourced 22 unique *T. thermophila* genotypes: 19 from the Cornell *Tetrahymena* Stock Center and 3 from the Chalker lab (Washington University, Appendix 1). These genotypes, which vary in geographic origin and have well-known genetic differences (Appendix 1), were chosen to sample existing genetic variation, not for their functional significance. As most were derived from laboratory cultures, our assemblage likely contains less variation than natural populations (Zufall *et al*. 2013), making our results a conservative estimate of how thermal adaptation might proceed in nature.

### Quantifying r, r-TPCs, and TPC shape parameters

Culture care and maintenance followed standard practices for ciliate Ecology (Appendix 2). Stock cultures were maintained in Percival AL-22 growth chambers with a 12hr day-night cycle at 22°C. We quantified *r*-TPCs for all genotypes through growth assays in 3cm diameter Petri dish microcosms containing 3mL of growth medium at seven temperatures (13, 19, 22, 25, 30, 32, 38°C). Each genotype and temperature combination was replicated six times, totaling 924 microcosms. Experimental temperatures span below and above the average growing season temperature in *T. thermophila*’s native range (∼23°C; NOAA 2024). Microcosms were initialized at densities of 1ind/mL in 3ml Petri dishes, which results in exponential growth for 1-2 days (Gibert *et al*. 2022, 2023; Singleton *et al*. 2021). After 24hrs, we censused the microcosms through whole-population counts under a stereomicroscope (Leica M205C) and calculated *r* across temperatures to characterize the entire *r*-TPC as log(final density/initial density)/time, with time = 1 day (Wieczynski *et al*. 2021). To obtain *r*-TPC shape parameters, we fitted a Sharpe-Schoolfield model (“nls.multstart” v1.3.0 package in R, (Padfield 2023)). Shape parameters E_a_, r_peak_, CT_min_ and T_opt_ (Fig 1b) could be unequivocally estimated from our data, and so we focused all subsequent analyses on those. These parameters control the rising portion of the *r*-TPC (Fig 1b, green) –the “operational temperature range” (DeLong *et al*. 2017), i.e., what most organisms are likely to experience in their native geographic ranges.

### Link between heritable genetic variation in r, and r-TPC shape parameters

We quantified E, G and G × E variation in *r* using function *gxeVarComps*() in R package statgenGxE v1.0.5. The function fits a linear model with *r* as the response variable, and temperature, genotype, and their interaction as fixed predictors, to calculate effect sizes and statistical significance. It then fits a second model with all fixed terms treated as random effects to calculate *r* variance components (E, G, and G × E). Only G is considered to be heritable, so the fraction of heritable genetic variation in *r*, can be estimated as the broad-sense heritability (H^2^=G/(E+G+G × E)). However, this expression does not account for inter-treatment and replicate variability, which inflates E and underestimates H^2^ (Cullis *et al*. 2006; Piepho & Möhring 2007). In the appendix we present three alternative formulations for H^2^ that address this issue (Appendix 3).

Next, we tested the predictions of our verbal model (Fig 1) linking heritable variation in *r* (G) to heritable variation in *r*-TPC shape parameters. Since each *r*-TPC yields a single shape parameter set, only G variation in shape parameters could be quantified. All other components of variation (e.g., E, G × E, error) were pooled as residual variance. To evaluate the model’s prediction that G variation in *r* results in greater G variation in shape parameters r_peak_ and CT_min_ compared to E_a_, and T_opt_ (Fig 1e), we tested whether r_peak_ and CT_min_ exhibited stronger genetic covariance with *r* than did E_a_, and T_opt_ using a multivariate Bayesian Generalized Linear Mixed Model. The model includes mean-scaled values of *r*, E_a_, r_peak_, CT_min_ and T_opt_ as jointly modeled response variables, each with their own fixed mean, and Genotype as a random effect to estimate variances and covariances implemented in R package MCMCglmm v.2.36 (Hadfield 2010).

Variance-covariance priors were obtained by bootstrapping *r*-TPC data 100 times within each genotype and re-fitting the Sharpe-Schoolfield model to each bootstrap replicate, generating a population of shape parameters for each genotype (Appendix 4). Using inverse Wishart priors (Murphy 2012) did not alter our results. Last, if G variation in *r* results in greater G in r_peak_ and CT_min_, these shape parameters should also have greater heritability than E_a_, and T_opt_. To confirm this, we calculated the broad-sense heritability in *r*-TPC parameters using the standard expression H^2^=G/(G+residual variance) with G and residual variance estimates from the MCMCglmm model.

### Consequences of heritable variation: selection and evolutionary potential of r-TPC parameters

Heritable *r*-TPC shape parameters can evolve under selection. To understand how, we quantified 1) the direction, form, and magnitude of selection on all four shape parameters, 2) whether and how temperature influenced selection, and, 3) their potential evolutionary responses across temperatures. We used two approaches: one that neglects genetic correlations between parameters but allows estimation of non-linear selection (e.g., stabilizing selection), and one that accounts for genetic correlations and neglects non-linear selection, while enabling predictions of evolutionary responses across temperatures.

In the first approach, we quantified selection by characterizing the adaptive landscape (Lande 1976, 1979) as the relationship between shape parameter values and absolute fitness—classically assumed to a function of *r* (Lande 1976, 1979) because it reflects average birth and death rates of individuals, two major fitness components (Lande 1982; Partridge & Harvey 1988). We modeled this relationship for all shape parameter values fitted across genotypes, using polynomial regression and *r* as the response, linear and quadratic effects (i.e., non-linear selection) for each mean-standardized shape parameter, and additive and interactive temperature effects with the linear and quadratic shape parameter terms (see Appendix 5). We multiplied the quadratic regression coefficient by two (Stinchcombe *et al*. 2008). A positive relationship between *r* and the focal shape parameter would be evidence of positive directional selection, a negative relationship would indicate negative directional selection, and no relationship suggests no directional selection. Stabilizing selection would result in a concave-down relationship, and disruptive selection would manifest as a concave-up relationship where extreme values have higher fitness (Lande & Arnold 1983).

The second approach predicts *r*-TPC shape evolution across temperatures while accounting for genetic correlations among shape parameters. It uses the multivariate breeder’s (Lande 1979) and Price’s (Price 1972) equations to estimate direct (i.e., acting on the focal parameter) and indirect selection (i.e., acting on genetically linked parameters, (Stinchcombe *et al*. 2014)). This is achieved by estimating the response to selection of all shape parameters, **Δz**, as their covariance with fitness—i.e., the Price equation aspect of this approach– which requires estimating a genetic variance-covariance matrix (**G_zw_**)—i.e., its multi-variate breeder’s equation aspect. This matrix includes the genetic variance-covariance matrix (**G**) of shape parameters as its first four rows/columns, and their genetic covariation with fitness (W) as the last row/column (Lande 1976, 1979, 1982). The finite rate of increase R=*exp(r)* was used as our metric of fitness but using plain *r* did not alter our results. From the Price equation, this last row/column also equals the vector of predicted trait change, **Δz**, so **G_zw_** also yields **Δz**. Then, from the multivariate Breeder’s equation, **Δz** = **Gβ**, where **β** is the selection gradient (Lande 1979). Provided that **G** is invertible, then **β** =**G**^-1^ **Δz**, to estimate selection.

Positive/negative Δz_i_ values indicate an increase/decrease in the i-th shape parameter, while positive/negative β_i_, indicates positive/negative selection operating on the i-th shape parameter. Alignment between Δz_i_ and β_i_ indicates direct responses to selection (Hansen & Houle 2008), in our case, by temperature, while misalignment suggest indirect selection through correlated responses with other shape parameters.

To estimate **G_zw_** for each temperature, we used subsets of *r* for each genotype for each temperature. Shape parameters do not change across temperatures but *r* does, so the **G_zw_** matrices only differed in their last row/column across temperatures, i.e., the covariances between shape parameters and fitness—or **Δz**. To facilitate this estimation and ensure interpretability, all **G_zw_** variables were rescaled, which was achieved through the expression **G_zw_** as **=SLS\***(1/2F) where **L** is a between-genotype covariance matrix, **S** is a diagonal matrix containing the inverse of each trait’s standard deviation, and F is the inbreeding coefficient, which equals 1 for clonal lineages (Falconer 1996). We quantified uncertainty in **G_zw_** entries using a Bayesian posterior distribution of **L** matrices (and hence **G_zw_**) and used a multivariate normal likelihood function and non-informative inverse regularized Wishart priors (nu=traits+1) (Murphy 2012), implemented in evolqg v3.0 R package (Melo *et al*. 2015). Using MCMCglmm instead of evolqg did not affect our results (Appendix 6). We took 1000 posterior samples for **G_zw_** to calculate **Δz**, **G**, and 𝛃, and 95% maximum density intervals for al estimates.

### Consequences of G × E in r for rapid r-TPC evolution

Because G × E variation in *r* is predicted to result in non-parallel *r*-TPCs (Fig 1), slight differences in intrinsic growth rates across temperatures can drive temperature-mediated differential selection among genotypes (Fig 4a, b). Treating *r* as proxy for absolute fitness (Lande 1976), this selection should be strongest at temperatures where *r*-TPCs differ the most, i.e., the temperatures with the largest differences in relative fitness.

We empirically tested for temperature-dependent selection among genotypes by setting up an experimental evolution assay with two fluorescently marked strains (AXS and CU4106, Fig 4c) with different *r*-TPCs and relative fitness (Fig 4d, Fig4d inset) across six temperatures (19, 22, 25, 30, 32, 38°C), each replicated seven times, and 2 additional single-strain controls per temperature. The genotypes cannot mate because they belong to the same mating type (mating type VII, Appendix 1), so only clonal reproduction was possible. We initialized our microcosms at equal densities (5 ind/mL in mixture treatments, 10ind/mL for single-genotype controls). After 48 hours, we added Cadmium Chloride to induce fluorescence (Appendix 7), confirmed on a Leica Thunder Cell Culture inverted microscope (Fig 4c and Appendix 8). Thirty minutes later, we cytometrically censused each sample (Novocyte 2000R). While fluorescently tagged genotypes can lose their ability to fluoresce over time, they carry a paromomycin resistance gene (Appendix 7), allowing selection through paromomycin exposure (100μg/mL) prior to counting and estimating relative frequencies based on fluorescence (Appendices 9-10). Antibiotics can negatively affect protists and the bacterial communities they feed on, so we replicated this experiment in paromomycin-free conditions, which did not qualitatively alter our results (see Appendix Fig S11).

Last, observed changes in genetic frequencies were compared to predicted frequencies by a classic model of genetic evolution in discrete time parameterized with the *r*-TPC of both experimental genotypes (Fig 4f). The model tracks the frequency of each strain 𝑓*_i_*, and assumes that their absolute fitness, 𝑊*_i_*, is a function of their average intrinsic growth rate *r* (Lande 1976, 1979). The frequency of each strain in the population is determined by the classic recursive equation, 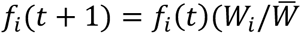) [Eq 1.], where 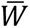 is the average fitness of the population (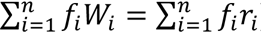) and 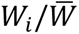 is fitness the relative fitness of i-th genotype *relative* to the entire population. We used each genotype’s *r*-TPC (AXS, CU4106, Fig 4d) to estimate 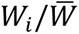 across temperatures (i.e., replacing 𝑟*_i_*). We forward-solved the recursive equation (Eq 1) under two scenarios: one in which temperature was constant, and one in which temperature fluctuated randomly across generations, drawn from a normal distribution with known mean and variance. This minimal model assumes that differential selection across environments is mostly driven by differences in *r*-TPCs, and disregards density- and frequency-dependent selection, sexual reproduction, mutation, and most all ecological processes.

## RESULTS

### Heritable variation in r leads to heritable variation in r-TPC shape parameters

Intrinsic growth rates showed strong temperature responses within genotypes (F = 3092.70, D_f_ = 6, Generalized Eta-Squared (GES) effect size = 0.964, p ≤ 0.001), significant variability across genotypes (F= 163.65, p ≤ 0.001, D_f_ = 21, GES = 0.832), and significant G × E interactions (F=29.7, p ≤ 0.001, D_f_ = 122, GES = 0.840), leading to classically unimodal shapes that varied among genotypes (Fig 2a). Environmental variation (E) accounted for 71.7% of all observed variation in *r*, genetic variation (G) explained 6.1% of all variation, and Gene-by-Environment interactions (G × E) explained 11.7%, with 10.5% residual variation (Fig 2b). This is well within what is expected for life history traits (Hoffmann & Sgrò 2011). Despite a relatively modest G compared to E, after accounting for experimental inter-treatment and replicate variability, *r* was strongly heritable (H^2^_standard_=0.76, H^2^_cullis_=0.95, H^2^_piepho_=0.91, Appendix 3).

**Figure 2.**
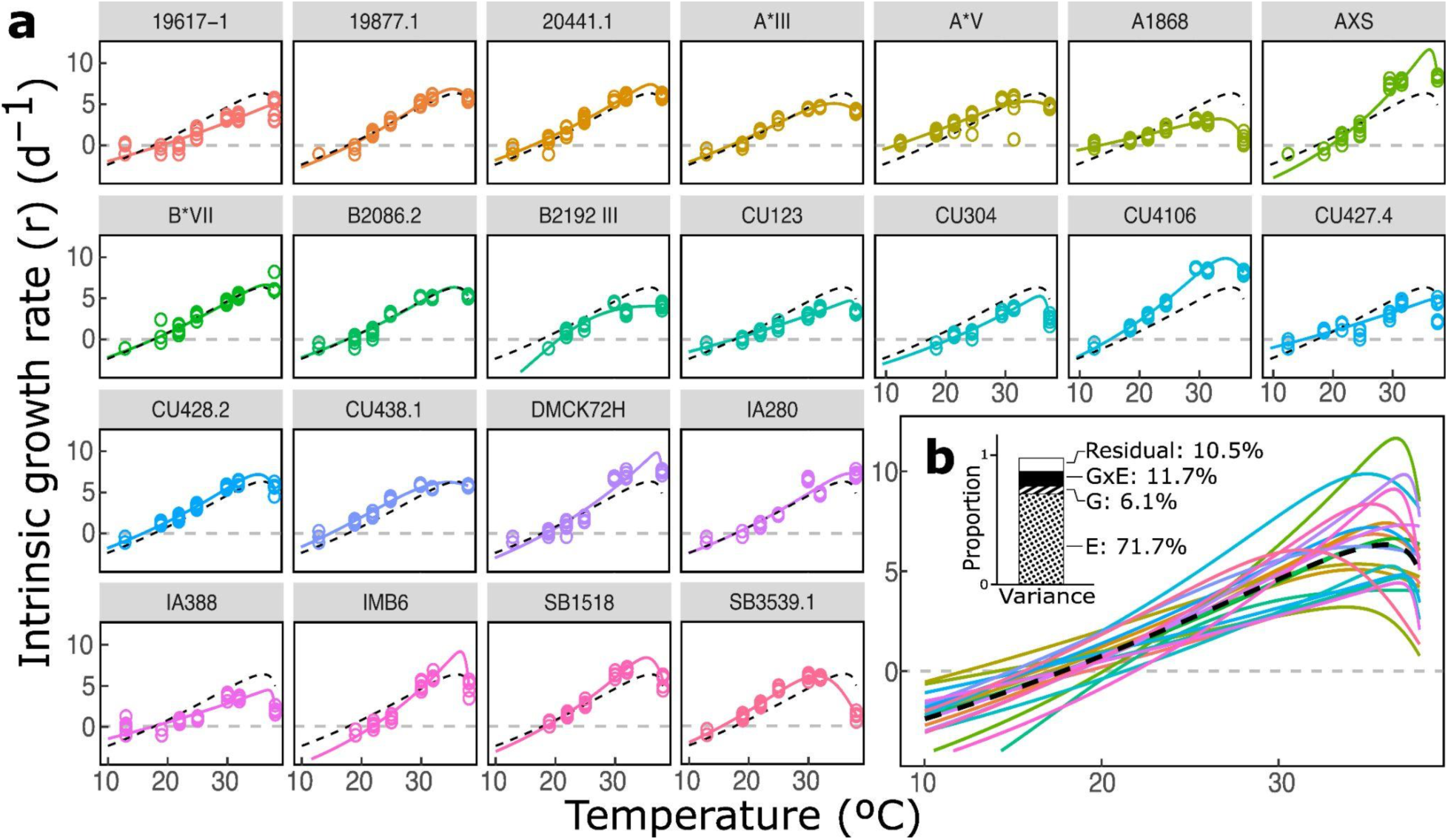
(a) Observed r-TPCs for all 22 genotypes. Dots represent observed *r* values, bold lines represent Sharpe-Schoolfield model fits, and dash lines represent average TPCs across all experimental genotypes. (b) All 22 TPCs are superimposed and dashed lines represent the average r-TPC. Inset: amount of variation due to residual, G × E, G, and E variation in *r*.

Shape parameters varied widely across genotypes (e.g., genotype AXS was 3.2 times more thermally sensitive than A1868 and could grow 3.6 times faster at T_opt_, Fig 2, and Appendices 12-13). As predicted, CT_min_ and r_peak_ showed strong genetic covariation with *r* (∣ 𝐺𝑐𝑜𝑣_CTmin_ ∣ = 0.65, ∣ 𝐺𝑐𝑜𝑣_rpeak_ ∣ = 0.63, Appendix 14), but there was no evidence of genetic covariation between *r* and T_opt_ or E_a_ (credible intervals contained 0, Appendix 14). Last, shape parameter heritabilities matched verbal model predictions (H^2^_CTmin_=0.945, H^2^_rpeak_=0.949, H^2^_Topt_=0.070, H^2^_Ea_=0.002), confirming that heritable variation in *r* results in heritable variation in CT_min_ and r_peak_, but not T_opt_ or E_a_.

### Consequences of G in r: selection and evolvability of r-TPC shape parameters

Without accounting for genetic covariances, selection operated differentially across shape parameters and was temperature dependent: r_peak_ was under negative directional selection at low temperatures (<20°C, Fig 3a, Appendix 15), weakly positive or no directional selection at intermediate temperatures (between 20 and 30°C, Fig 3a, Appendix 15), and strong positive directional selection in high temperatures with E_a_ following a similar pattern (Fig 3b, Appendix 16). CT_min_ was under negative selection at low/intermediate temperatures but no selection at high temperatures (Fig 3c, Appendix 17). Last, T_opt_ was under no selection at low temperatures but under weak then strong stabilizing selection at intermediate and high temperatures, respectively (Fig 3d, and Appendices 18-19).

**Figure 3.**
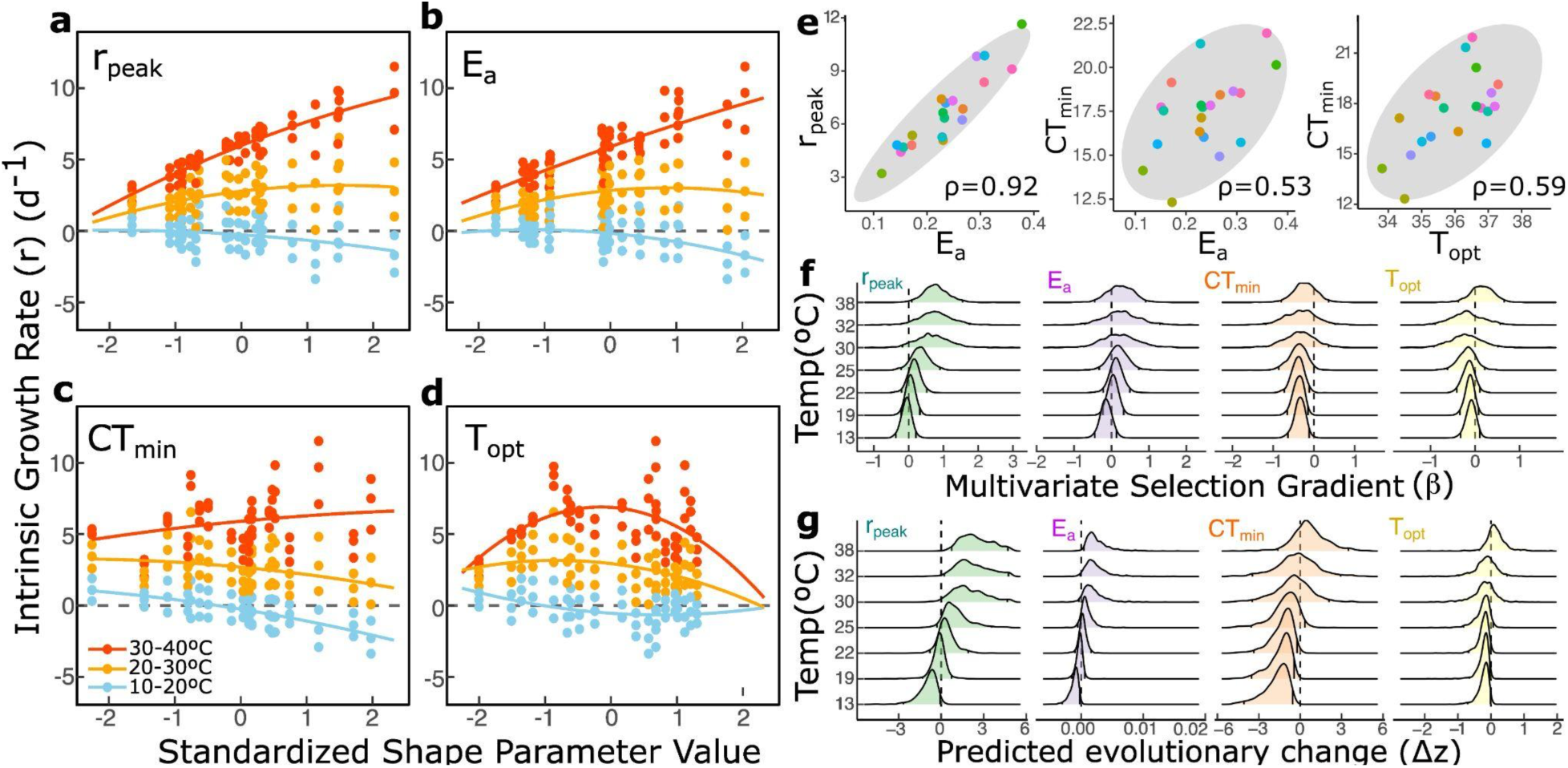
(a) Estimated adaptive landscape across temperatures (i.e., change in fitness with a change in the underlying shape parameter) for r_peak_. Color indicates temperature bins (blue: 10— 20°C, yellow: 20—30°C, red: 30—40°C) (b) As in a, but for E_a_. (c) As in a, but for CT_min_. (d) As in a, but for T_opt_. (e) Observed genetic associations between shape parameters. Each dot is a genotype color-coded as in Fig. 2. In gray, 95% confidence ellipses. ρ represents correlation coefficients. (f) Estimated multivariate selection coefficient (β, 95% maximum density intervals) for all shape parameters across temperatures, or eco-evo landscapes(MacColl 2011). (g) Predicted evolutionary change (Δz, 95% maximum density intervals) for all shape parameters, across temperatures.

However, we also found clear positive genetic covariances between r_peak_ and E_a_, CT_min_ and E_a_, and CT_min_ and T_opt_ (Fig 3e). Consequently, selection acting on highly heritable parameters CT_min_ and r_peak_ could still result in correlated evolution in weakly heritable parameters E_a_, and T_opt_. Accounting for these genetic covariances, we confirmed differences in selection across shape parameters whose magnitude and direction also shifted with temperature (Fig 3f), resulting in possible temperature-dependent shifts in parameters (Fig 3g). Specifically, our multivariate analysis suggested that selection would favor higher r_peak_ (and maybe E_a_) at high temperatures, and low E_a_ and CT_min_ at low temperatures, but no selection on T_opt_ (Fig 3f), generally matching univariate predictions (Fig 3g). Overall, the evolutionary responses followed predicted trajectories from the estimates of selection closely (Fig 3f-g), with a few exceptions that suggest correlated evolution: positive response of E_a_ with temperature likely are correlated responses with r_peak_ (Fig 3g), which in turn shows strong negative response at low temperatures despite no selection, likely through correlated evolution with E_a_ whose negative response is driven by negative selection in CT_min_ (Fig 3g). Despite differences in heritability which impose constraints in r-TPC evolution, genetic correlations among shape parameters allowed for temperature-dependent evolutionary responses in all of them except T_opt_, which our results suggest is under stabilizing selection (Fig 3d).

### Consequences of G × E in r: sorting of standing genetic variation across temperatures

We observed significant temperature-dependent differential selection among genotypes (Fig 4e), matching theoretical predictions from a simple model of genetic evolution (Fig 4f), which confirms the hypothesis that G × E in *r* could set the stage for rapid r-TPC shape evolution through selection on standing genetic variation (Fig 4a, b). Despite quantitative discrepancies between observed and predicted frequencies—notably at 19°C where the model predicted a polymorphic population but the data indicated otherwise (Fig 4e, f)—it still correctly predicted observed changes in genetic frequencies across most temperatures. These results are also robust to different experimental conditions (Appendix 11) and fluctuating temperatures (Appendix 20), the latter showing that temperature fluctuations could play a role in determining r-TPC evolution if *r*-TPCs show multiple crossing points in a short temperature span.

**Figure 4.**
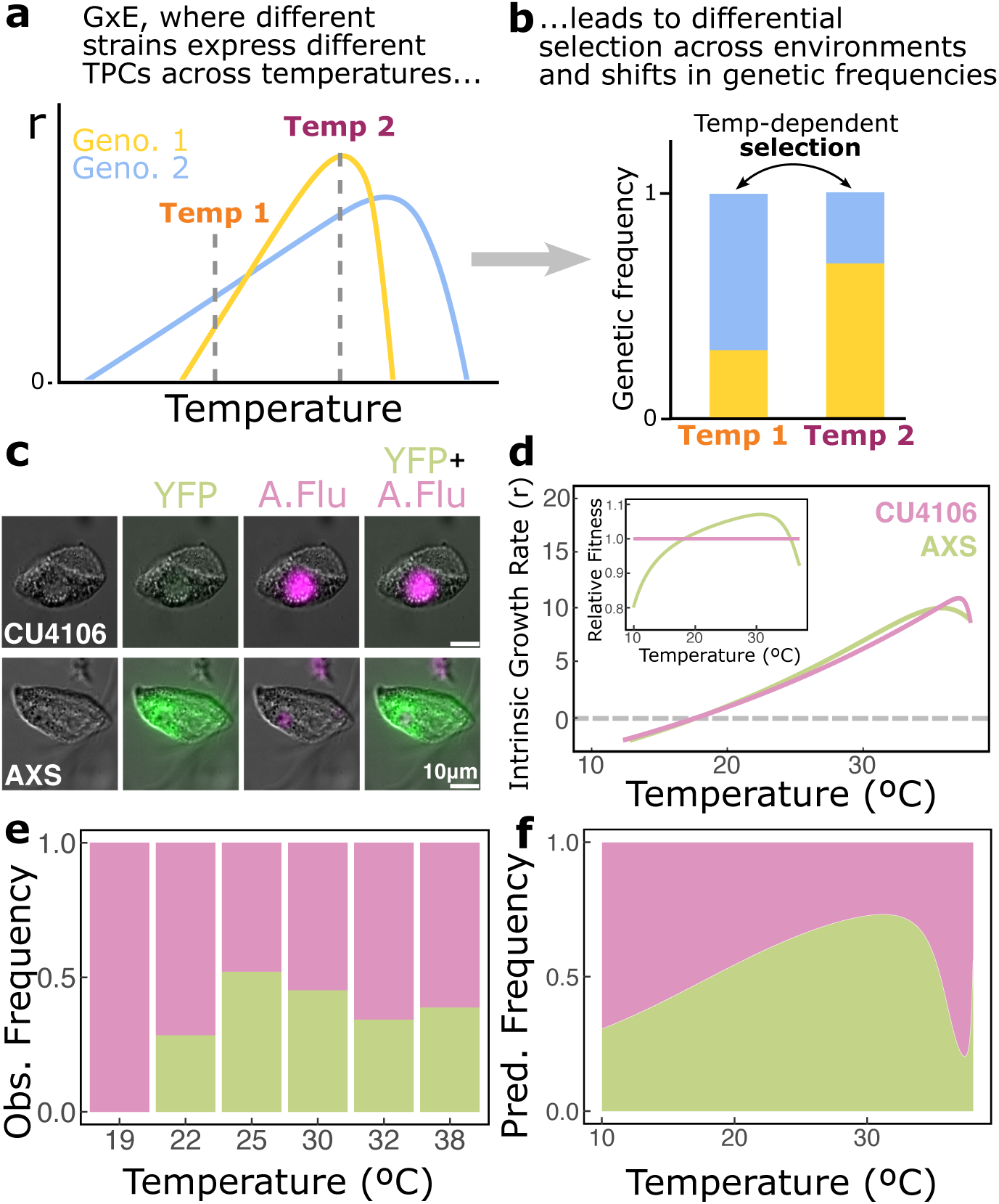
(a) G × E variation in r-TPCs leads to differential growth of each genotype across temperatures. (b) Differential growth across temperatures leads to differential selection across environments and rapid shifts in genetic frequencies across temperatures. (c) First column: Differential Interference Contrast (DIC) microscopy for two genotypes of the protist *Tetrahymena thermophila* (CU4106 and AXS). Second column: fluorescence microscopy image overlayed on DIC. Only AXS fluoresces (green) due to the expression of Yellow Fluorescent Protein (YFP). Third column: as in the second column, but for autofluorescence (A. Flu, in pink), which both genotypes exhibit. Fourth column: Overlayed DIC, YFP and A.Flu images showing how the different strains fluoresce once all sources of fluorescence are accounted for. (d) r-TPC for genotypes CU4106 and AXS. Inset: Measures of relative fitness for both CU4106 and AXS. This predicts an increase in AXS frequency relative to CU4106 at intermediate temperatures relative to low or high temperatures. (e) Observed genetic frequencies across temperatures. (f) Predicted genetic frequencies across temperatures.

The results of this experiment were also consistent with our estimated predicted responses to selection (cf. Fig 3 and Fig 4): lower temperatures led to higher frequencies of the CU4106 genotype, whose *r*-TPC has lower E_a_ and CT_min_ compared to AXS (Fig 4d), so the average *r*-TPC of the population should reflect that and also have lower E_a_ and CT_min_, as predicted by our multivariate model. At higher temperatures, selection favored AXS, which has higher E_a_, so the ensuing population also should have an average *r*-TPC with higher E_a_ (Fig 4d). Last, rising temperatures led to an increase in genetic variance (∼p(1-p) where p is the frequency of either genotype and 1-p that of the other) compared to lower temperatures (Fig 4e, f)—which should facilitate adaptation—then a reduction in genetic variance (as genotype CU4106 becomes less prevalent, Fig 4e, f), which in turn could impede adaptive evolution in the future.

## DISCUSSION

Our study reveals heritable genetic variation in *Tetrahymena thermophila*’s population intrinsic growth rates that results in heritable variation in some, but not all, *r*-TPC shape parameters (Fig 2), allowing for thermal adaptation in new climates (Fig 3). Observed *intra*-specific variation in *r*-TPC shape parameters rivals reported *inter*-specific variation across multiple protist species (Wieczynski *et al*. 2021). We also found that most *r*-TPC shape parameters are under different selection regimes across temperatures (Fig 3), with parameters CT_min_ and r_peak_ being more likely to respond to selection than E_a_ and T_opt_, but even weakly-heritable parameters can still evolve through correlated evolution (Fig 3). Consequently, colder temperatures select for lower CT_min_, r_peak_, and E_a_, and warmer temperatures lead to high r_peak_ and E_a_ (Fig 3). Last, we showed that G × E in *r* can lead to rapid, predictable shifts in population genetic makeup across temperatures and *r*-TPC evolution (Fig 4), in a “plasticity drives adaptation” scenario (Ghalambor *et al*. 2007).

While the evolution of microbial TPCs is likely the product of adaptation to local habitats (Kontopoulos *et al*. 2020), how *r*-TPCs will adapt to ongoing rising temperatures remains an open question. Several canonical evolutionary paths have been proposed: 1)‘Colder-is-Better’ (CIB), where rising temperatures reduce population growth, leading to lower r_peak_, T_opt_, and E_a_ (Kingsolver & Huey 2008). 2) Warmer-Is-Better (WIB), where higher growth rates evolve in warmer temperatures, leading to TPCs with higher r_peak_ and T_opt_ (Pawar *et al*. 2015). And, 3) Generalist-Specialist-Tradeoff (GST), where species can either evolve towards rapid growth within a narrow temperature range (i.e., temperature specialists), or slower growth over a broader temperature range (i.e., temperature generalists), leading to higher r_peak_, higher CT_min_, and lower CT_max_ (Seebacher *et al*. 2015). Tests of these possible evolutionary paths (Kontopoulos *et al*. 2020; Montagnes *et al*. 2022) mostly use inter-species comparisons that often overlook intra-specific variation and genetic associations between shape parameters, and therefore cannot readily make predictions about *r*-TPC evolutionary trajectories for any given species. Indeed, without accounting for genetic associations, our results would suggest support for WIB with clear directional selection for higher r_peak_ and E_a_ under warming climates (Fig 3a, b). Accounting for genetic covariances, however, suggested more complex evolutionary *r*-TPC responses than currently predicted by theory. Specifically: we show support for WIB as warming should favor *r*-TPCs with high r_peak_ and high E_a_ (Fig 3f, g), but no support for GST, as CT_min_ and r_peak_ responded mostly together (Fig 3f, g). Last, T_opt_ is assumed to evolve under WIB, CIB and GST, but we found no selection (Fig 3f) –or maybe even weakly stabilizing selection (Fig 3d)– and no evolutionary response (Fig 3g). This suggesting that neither WIB, CIB and GST can explain predicted evolutionary responses in *T. thermophila*, which emphasize the importance of intraspecific variation and genetic associations to fully understand *r*-TPC evolution.

When G × E in *r* is prevalent, we showed that thermal adaptation can occur rapidly through temperature-dependent selection on *r*-TPC genotypes (Fig 4), resulting in reduced genetic variation. Loss of G which could slow down, or impede, future adaptation (Pauls *et al*. 2013). In other words, we show that phenotypic plasticity in *r* could lead to adaptive evolution in r-TPCs in novel climates. That this form of r-TPC evolution could be anticipated from simple models of genetic evolution, is striking, considering how little information beyond the r-TPC of each genotype the model accounts for. Yet, *r*-TPC adaptation in deep time has been suggested to occur gradually across six different *Tetrahymena* species (Montagnes *et al*. 2022), directly countering our claim that *r*-TPCs could evolve rapidly through temperature-dependent selection. We argue that this form of rapid *r*-TPC evolution may only play out during fast-changing environmental conditions, not always result in long-term *r*-TPC change, or be masked by reversals in selection. In fact, selection often “erases its traces” (Haller & Hendry 2014), and is strongest under novel conditions, weakening as populations adapt (Caruso *et al*. 2017)—i.e., “the paradox of stasis” (Haller & Hendry 2014).

Resulting temperature-dependent loss of genetic diversity of adaptive r-TPC evolution could impact the persistence of genetically depauperate species under warming. But whether this may be the case in nature hinges on quantifying processes that generate diversity (mutation, gene flow, recombination) in *r* and *r*-TPC shape parameters in the wild, and assessing when these processes could counter loss of genetic diversity after episodes of thermally-induced *r*-TPC evolution. *T. thermophila* has the lowest reported base substitution mutation rate of all species (Zufall et al. 2016), and limited gene flow (Zufall et al. 2013), but even weak dispersal or rare mutation events could contribute to *r* and *r*-TPC variation and rescue genetic diversity in very large, rapidly reproducing populations. The prevalence of sexual reproduction in nature is debated –e.g. ∼50% of wild individual *T. thermophila* cannot reproduce sexually and some *Tetrahymena* lineages have lost this ability entirety (Doerder 2014)– but likely important if present. In fact, *T. thermophila* possesses two separate genomes, a germinal line (in micronucleus) and an expressed genome (in macronucleus). Our study focuses on macronuclear variation, but asexual reproduction following sex results in random distribution of macronuclear alleles to daughter cells—i.e. phenotypic assortment—potentially generating novel variation in r and r-TPCs (Tarkington *et al*. 2023). Moreover, seemingly lost genetic variation could persist in the population in other individual’s micronuclei and reemerge following sexual reproduction (Dimond & Zufall 2016). Fully understanding thermal adaptation in nature thus requires exploring these processes’ interplay, yet, for most organisms, they remain poorly understood.

### Caveats

While bodies of water in *T. thermophila*’s native range are warming significantly (US EPA 2021) and experiencing more frequent and extreme heatwaves (Tassone *et al*. 2023), water temperature typically remains 2-5°C cooler than air temperature (Stefan & Preud’homme 1993). Thus, some of the warmer temperatures in our study may be uncommon in *T. thermophila*’s native range. However, strong temperature effects observed at low temperatures suggest these patterns can still operate in many *T. thermophila* populations. Also, previous research showed that most variation in *r*-TPC shape across species could be collapsed into a single dimension of variation (Rezende & Bozinovic 2019), suggesting that some of the covariation observed in this study between shape parameters could be spurious. If these *r*-TPCs could actually be collapsed into a single axis of variation, however, we would have expected strong and multiple covariation between most if not all shape parameters—which was not the case (Fig 3e, f). Interestingly, most of the genetic variation in shape parameters was observed, as predicted, in CT_min_ and r_peak_, but these two parameters were not correlated to one another. Perhaps our intraspecific *r*-TPC variation results stand in contrast to those observed across species (Rezende and Bozinovic, 2019), but more research is needed to clarify this point.

### Conclusion

Overall, TPCs control the fate of populations (Sinclair *et al*. 2016), ecological interactions (Enquist *et al*. 2015), food web dynamics (Barbour & Gibert 2021; Gibert *et al*. 2022), and ecosystem processes (Gibert *et al*. 2015). Yet, TPC evolution in a rapidly warming world remains conspicuously unknown. Here, we show that *r* intraspecific variation can drive temperature-dependent evolution in microbial *r*-TPCs and have consequences for rapid shifts in population genetic makeup. Despite studying a single protist species, our findings provide a foundation for studying thermal adaptation in this diverse and important microbial group and beyond. Our study emphasizes the importance of temperature in mediating rapid microbial evolutionary change as we grapple with understanding and predicting organismal responses to an increasingly warmer world.

## STATEMENT OF AUTHORSHIP

All authors contributed to the conceptual foundation of the study. MHL, ZH, and JPG designed the study. DC contributed the fluorescent lines and provided cell biology expertise along with MO. MHL, ZH and AY collected the data. MHL and JPG performed data analyses with support from DHW, RZ, and AS. JPG did the mathematical modeling and FAM the quantitative genetics. MHL and JPG wrote the first draft of the manuscript and all authors contributed significantly to revisions.

## DATA ACCESSABILITY STATEMENT

All software and data from this study are available on GitHub (https://github.com/JPGibert/TPC_evolution).

## ACKNOWLEDGMENTS

We are grateful to Enzo Bruscato and Heather Raslan for their data collection support. We are indebted to Samraat Pawar, John Stinchcombe, and two anonymous reviewers for their feedback on earlier versions of our manuscript. AMS was supported by NSF DEB award 2306183. DLC acknowledges grant support for the Tetrahymena Stock Center by NIH project # 2P40OD010964 and additional support from NSF award # MCB 1947526. JPG acknowledges generous support from DOE BER DE-SC0020362 (which also supported MHL, AY, and DJW), NSF DEB award number 2224819, NSF CAREER award number 2337107, and a Simons Foundation Early Career Fellowship in Microbial Ecology and Evolution LS-ECIAMEE-00001588 (which also supported MHL and AY).

