## Supplementary_Appendix for "Rapid adaptive evolution of microbial thermal performance curves"

Online Supplementary Materials: Rapid adaptive evolution of microbial thermal performance curves

INDEX:

**Appendix 1. Genotype Table**

Table S1: Table of genotype data and metadata.

| Name | Original Provenance | Provenance | Stock ID | Mutations |
|---|---|---|---|---|
| 20395-1 | Lake Warren in Alstead, NH, lat. 43 07.310, long. -72 17.840 | Cornell Tetrahymena Stock Center | SD01557 | NA |
| SB3539-I | | Cornell Tetrahymena Stock Center | SD00660 | *chx1[C3]-1/chx1[C3]-1 (CHX1[C3]; cy-s, I)*,<br><br>C3 strain - functional heterokaryon carrying cycloheximide resistance in micronucleus. |
| B*VII | | Cornell Tetrahymena Stock Center | SD00023 | B strain star line. Lacks a genetically functional micronucleus. |
| B2192 III | Frankel lab (Leslie Jenkins) | Cornell Tetrahymena Stock Center | SD01754 | Derived from a cross of B2086 II x B2086 VIa. Isogenic with B2192 IVB |
| CU428.2 | | Cornell Tetrahymena Stock Center | SD00178 | *mpr1-1/mpr1-1* (*MPR1*; mp-s, VII) |
| DMCK72 H | Chalker Lab, Washington University in St. Louis | | | |
| 19877 | SG69-4 in Guys Mills, PA (lat. 41 38.023, long. -79 53.514, elevation 1660 ft) | Cornell Tetrahymena Stock Center | SD01555 | Cech's self-splicing intron is present. Cytochrome oxidase I haplotype = WPA1 |
| 19617-1 | FS136E in PA (latitude 41.46, longitude -78.88) | Cornell Tetrahymena Stock Center | SD03089 | Has a micronucleus. No mating, immature. |
| IMB6 (GFP) | Chalker Lab, Washington University in St. Louis | | | |
| A*V | | Cornell Tetrahymena Stock Center | SD00014 | Lacks a genetically functional micronucleus. |

| | | | | |
|---|---|---|---|---|
| 20441-1 | Gregg Lake in Antrim, NH (lat. 43 02.605, long. -71 59.383 | Cornell Tetrahymena Stock Center | SD01560 | |
| CU427-4 | | Cornell Tetrahymena Stock Center | SD00715 | chx1-1/chx1-1 (CHX1; cy-s, VI) |
| SB1518 | | Cornell Tetrahymena Stock Center | SD01537 | *gal1-1/gal1-1; tyr-14/tyr-14* |
| IA388 | | Cornell Tetrahymena Stock Center | SD01454 | *elo1-1/elo1-1* (*elo1*; II) |
| 21157-1 | FS343S in PA (latitude 41.45, longitude -78.88) | Cornell Tetrahymena Stock Center | SD03114 | |
| CU438-1 | | Cornell Tetrahymena Stock Center | SD00189 | *pmr1-1/pmr1-1* |
| CU304 | | Cornell Tetrahymena Stock Center | SD00051 | CHX1/CHX1; chx2-1/chx2-1; mpr1-1/mpr1-1 |
| CU4106 | | Cornell Tetrahymena Stock Center | SD01010 | *mpr1-1/mpr1-1 Nulli 4 (MPR1; mp-s, VII)* |
| AXS | Chalker Lab, Washington University in St. Louis | | | YFP fusion into the BTU1 locus of Tetrahymena Strain CU428.2 (RRID TSC SD00178) |
| C*III | | Cornell Tetrahymena Stock Center | SD00024 | C strain star line. Lacks a genetically functional micronucleus. |

### Appendix 2. Culture Care

Upon reception, we transferred the cultures from axenic Proteose Peptone growth medium to Timothy Hay growth medium inoculated with a bacterial community from Duke Forest Gate 9 pond/Wilbur pond (Lat 36.013914, Long -78.979720, fully described elsewhere (Rocca *et al.* 2022)) and a wheat kernel as a Carbon source (Altermatt *et al.* 2015). We maintained these stock cultures in Percival (Perry, IA) AL-22 growth chambers under light (12hr day-night cycle) and temperature-controlled conditions (22°C) in 250mL borosilicate jars filled with 150mL of liquid medium refreshed every two weeks with new medium. As *T. thermophila* is a natural bacterivore, these experimental conditions are more realistic than purely axenic ones.

**Appendix 3. Estimating *r* heritability ($H^2$)**

We estimated E, G and G × E of *r* using the function *gxeVarComps*() in R package statgenGxE v1.0.5. The procedure fits two models: first, it fits a fixed effects linear model with *r* as the response variable, and temperature, genotype, and the interaction between temperature and genotype as predictors to calculate effect sizes, significance levels, and Best Linear Unbiased Estimators (BLUEs, (Baksalary & Puntanen 1990; Henderson 1975)). BLUEs are subsequently used to calculate *r* broad-sense heritability (see below). Second, it re-fits the model with all terms as random effects to calculate the variance component of each term (i.e., G, E and G × E) and Best Linear Unbiased Predictors (BLUPs, (Henderson 1975)), which are also subsequently used in the calculation of heritability. Using function *H2cal*() in R package inti v0.6.2 we calculated the broad-sense heritability ($H^2$) in three ways: 1) standard heritability, where$H^2$=G/P, P = G+(G×E/m)+(ResidVar/(m*r)), *m* is the number of temperature treatments and *r* the number of replicates (which accounts for inter-treatment and replicate variability , in ways that G/P does not (Baksalary & Puntanen 1990; Henderson 1975), 2) *Cullis* heritability, where $H^2$= 1-vBLUE/2G, and vBLUE is the mean variance of the difference of two BLUEs(Cullis *et al.* 2006) and, 3) *Piepho* heritability, where $H^2$= G/(G+vBLUP), and vBLUP is the mean variance of a difference of two BLUPs (Piepho & Möhring 2007).

These approaches to calculate *r* heritability account for the propagation of errors that are a consequence of repeatedly measuring *r* across and within genotypes and temperatures. This compounded error results in replicate and inter-treatment variability that a naïve statistical modeling approach would be unable to properly model, resulting in higher-than-expected residual variation, thus deflating G relative to all other components of P. This in turn results in an underestimation of actual G, and hence, $H^2$.

### Appendix 4. Bootstrapped TPC Curves

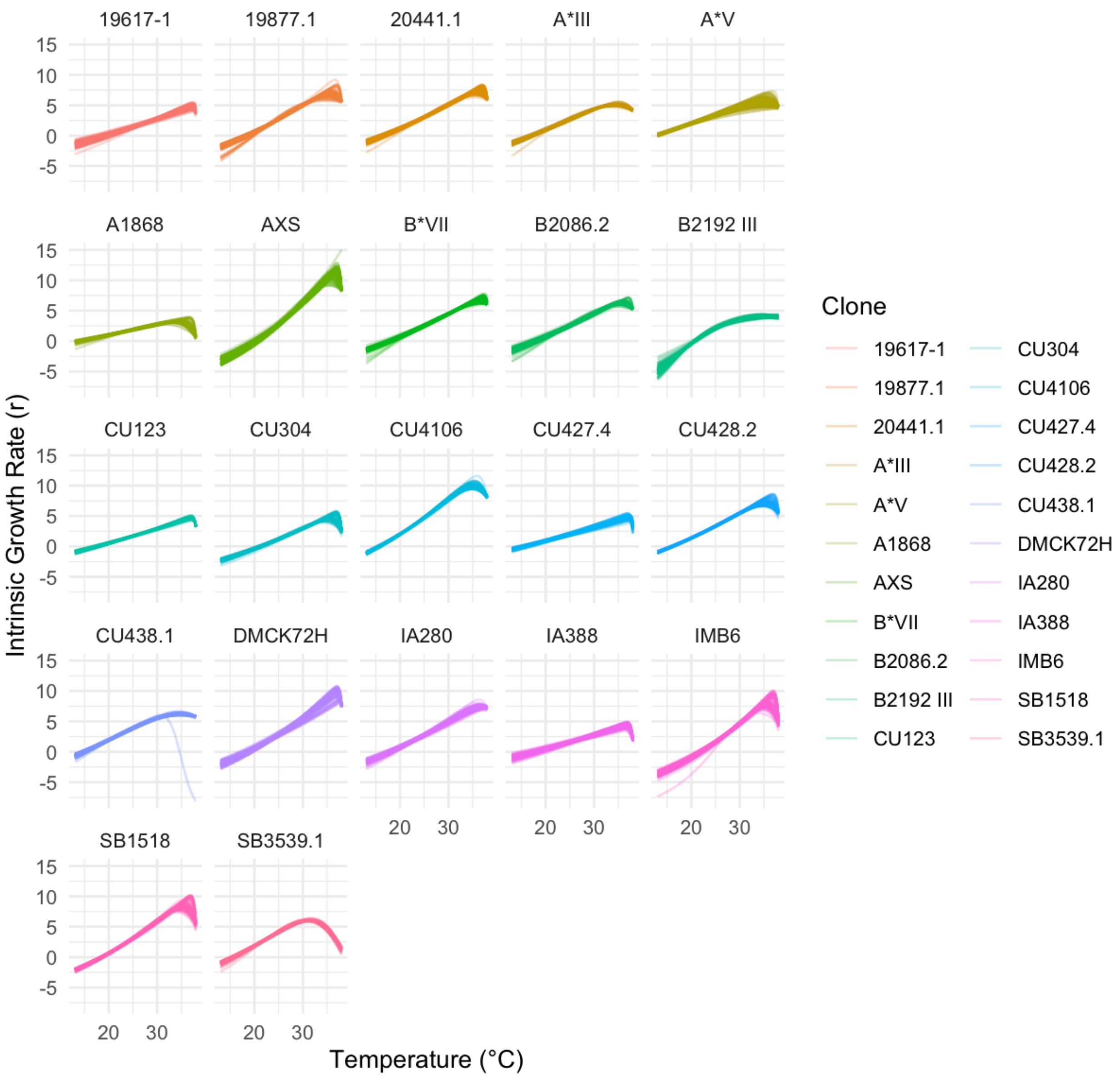

**Appendix 4 Figure S1:** Bootstrapped model fits across genotypes.

**Appendix 5. Quantification of the *r* adaptive landscape.**

Table S1: Model formulation for each shape parameter.

| Shape parameter | Model |
|---|---|
| $CT_{min}$ | $r \sim CT_{min}*Temp + CT_{min}{}^2*Temp$ |
| $E_a$ | $r \sim E_a*Temp + E_a{}^2*Temp$ |
| $r_{peak}$ | $r \sim r_{peak}*Temp + r_{peak}{}^2*Temp$ |
| $T_{opt}$ | $r \sim T_{opt}*Temp + T_{opt}{}^2*Temp$ |

**Appendix 6: Comparison Between evolqg and MCMCglmm estimates.**

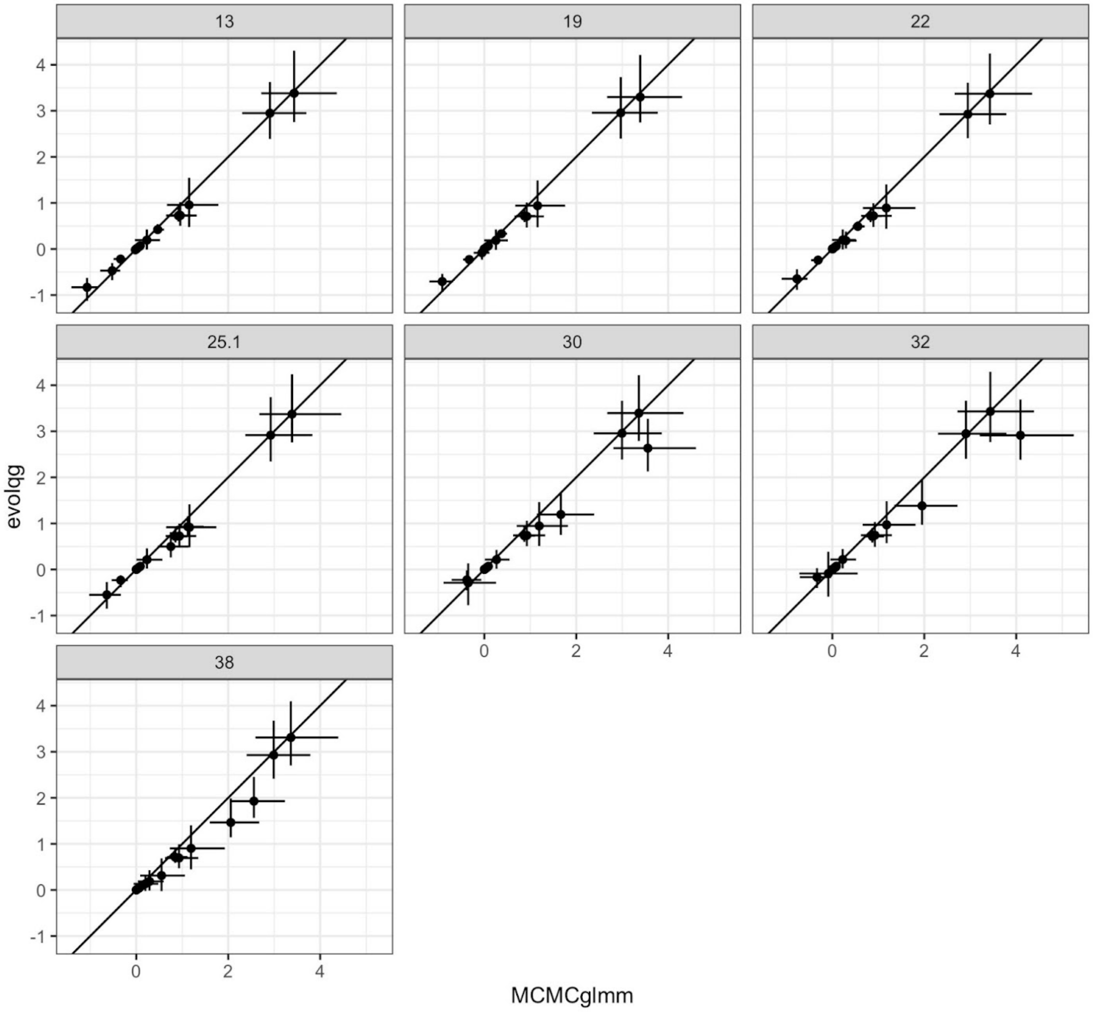

**Appendix 6 Figure S1:** Comparison between matrix parameter estimates (variances and covariances) between the evolqg function CalculateBayesianMatrix and the result from the MCMCglmm package. Each panel represents estimates for each temperature. Dots are median estimates and horizontal and vertical lines represent the interquartile range of the posterior distribution, with both approaches yielding the same result.

### Appendix 7. Methods For Creation of Fluorescent Strains and Fluorescence Induction

The AXS-YFP strain was generated by integrating a cadmium inducible TTHERM_00554360-YFP fusion into the BTU1 locus of Tetrahymena Strain CU428.2 (RRID TSC_SD00178) as described in Cole *et al.* 2023. Briefly, the coding region of Tetrahymena gene TTHERM_00554360 was amplified from Tetrahymena genomic isolated from strain SB210 (RRID TSC_SD01539) by PCR using Phusion (New England Biolabs, Ipswich NY) and two oligonucleotide primers: 5'-5'-CACCTTTAAAATGGCCACAAAAAAGTAGTGT-3' and GGATCCTTTATTTTTCTTGCCTTTTTTACC-3'. The purified product was first clones into plasmid pENTR before recombination into the gateway-compatible destination vector pBTUN5-ICYgtw by using LR clonase II. This expression plasmid was then introduced into Tetrahymena cells by biolistics transformation (Cassidy-Hanley *et al.* 1997). Plasmids contained a paromomycin-resistance gene, which allow transformants to be selected by growth in paromomycin containing medium. The creation of this strain was first reported on suprDB, an online research report database of unpublish results primarily generated in undergraduate classrooms (SuprDB 2021).

We started our experiment on Day 1, and initialized our microcosms at equal densities (5 ind/mL per genotype, and 10ind/mL for the single-genotype controls). Exactly 48hrs later (Day 3), we added 0.1μL of a 1mg/mL Cadmium Chloride ($CdCl_2$) solution to a 100 μL volume sample of each microcosm to reach a final concentration of 1 μg/μL of $CdCl_2$ in the sample, to induce fluorescence (Cole *et al.* 2023). After 30 minutes, we destructively harvested the microcosms to census them via cytometry. Between Days 1 and 2 the populations are growing exponentially, and density dependence is setting in by Day 3 –albeit still displaying extremely fast growth. It will take another 1.5-2 days for Tetrahymena to reach carrying capacity (Gibert *et al.* 2022)

### Appendix 8. Methods For Differential Fluorescence Imaging Across Strains

To confirm fluorescence of the two strains (Fig 4c), cells were mounted on glass slides with mounting medium (Winey *et al.* 2012). Images were taken with a Leica Thunder Cell Culture inverted microscope equipped with an HC PL APO 63X/1.40 N.A. oil-immersion objective lens. Fluorescent signals were captured using a 510-nm excitation laser and a 535/15-nm emission filter for YFP (expressed in genotype AXS), and a 395-nm excitation laser and a Leica DFT51011 quad-band filter set for autofluorescence (exhibited by both genotypes). All images were captured at a single Z plane using the same exposure settings; the resulting images were processed in ImageJ (see above and Fig 4c).

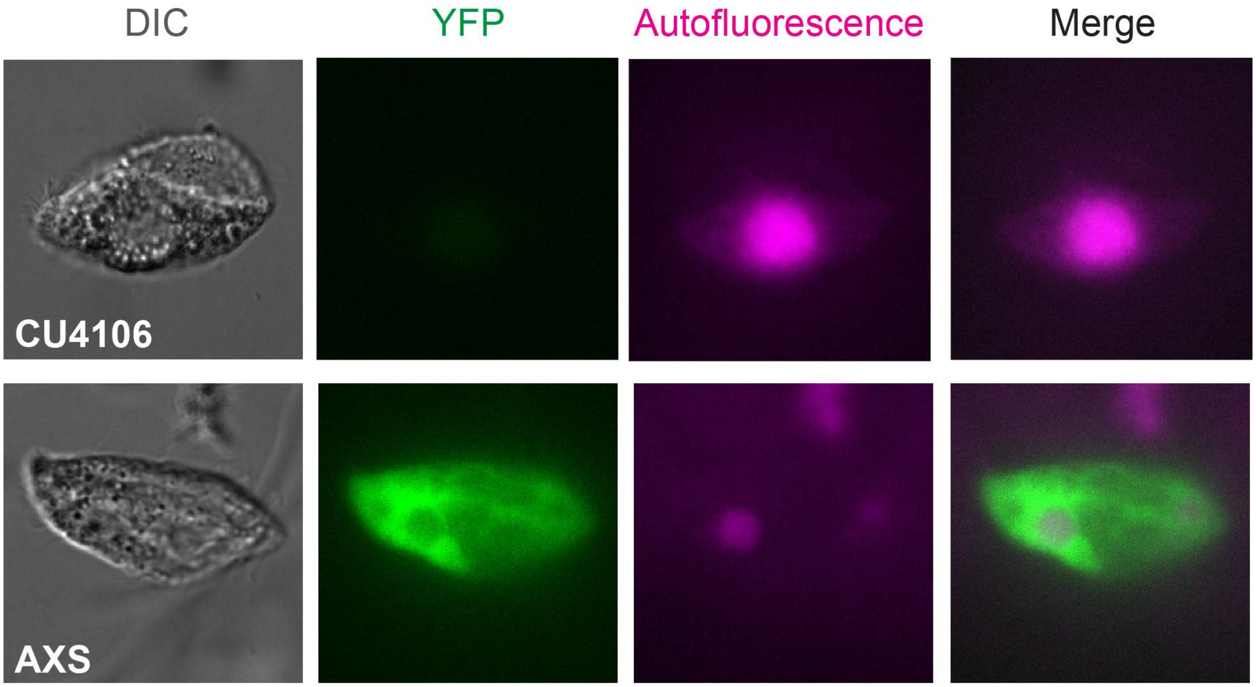

**Appendix 8 Figure S1:** 1st column: Differential Interference Contrast (DIC) microscopy for CU4016 and AXS protist *Tetrahymena thermophila* (CU4106 and AXS). Subsequent columns display raw fluorescence microscopy images. Photos are unimposed and uncorrected for relative fluorescence levels.

**Appendix 9: Flow Cytometry Frequency Calculations**

Each microcosm was censused with a Novocyte 2000R flow cytometer and analyzed using NovoExpress software v15.0. The flow cytometer detects particles (e.g., cells, debris, bacteria) based on how they scatter light and fluoresce (McKinnon 2018). Light scattering properties can be used to quantify cell size (FSC-H), and we detected fluorescence in the Phycoerythrin (PE-H, yellow) and Fluorescein isothiocyanate (FITC-H) channels. We gated the data in NovoExpress to select for the largest particles which in our case were all Tetrahymena *thermophila* cells. The data are plotted in Appendix S9 Fig 1. We used a PE-H versus FITC-H plot to parse the different fluorescent signals between the two experimental strains: CU4106 (which autofluoresces exclusively) and AXS (which autofluoresces and expresses Yellow Fluorescent Protein, or YFP). Control microcosms, which contained exclusively one of either strain for each temperature treatment, were used to determine the exact expected fluorescence range for each individual cell. An "AutoF" and "YFP" gate were created based on these controls (Figure 1). These control gating filters were then applied over each experimental microcosm, allowing us to identify cell strain based on their fluorescence pattern.

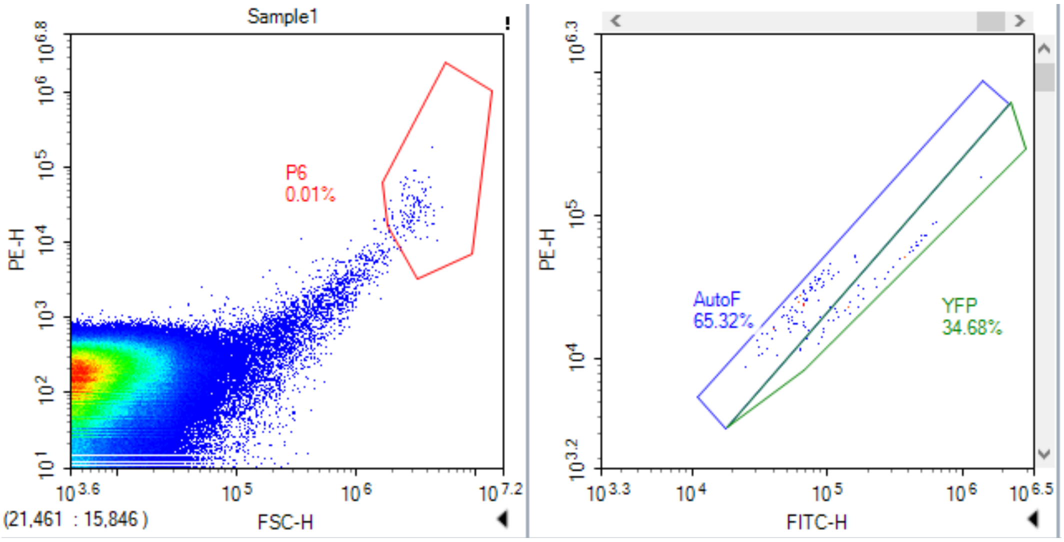

**Appendix 9 Figure S1:** This plot shows an AXS control sample, where 65% of AXS individuals fluoresced in the autofluorescence gating range.

We used two CU4106 control microcosms and nine AXS control microcosms per temperature. Additional AXS controls were necessary to increase detection precision while gating. Non-YFP tagged cells (CU-4106) fluoresced more weakly in the FITC-H channel than YFP-tagged cells (AXS) and generally fluoresced more strongly in the PE-H channel than non-YPF tagged cells fluorescence. CU4106 controls were detected exclusively in the autofluorescent gate. However, AXS controls were detected in both the autofluorescent (“AutoF”) and YFP gates, meaning that we could expect AXS cells to show up in the AutoF gate under experimental conditions, thus making it harder to parse YFP-tagged from non-YFP tagged cells.

To resolve that, we thus used the control microcosms to adjust the relative frequencies of each strain in each experimental microcosm, calculated as follows. For each single-strain control microcosm, we calculated the proportion of cells detected in each of the two gates across temperatures (Appendix S10). We then used the proportion of AXS cells across control replicates to adjust the observed number of AXS cells (i.e., cells showing up on the YFP gate) to ensure our estimate was as accurate as possible. We then subtracted this adjusted count from the total number of individuals in each microcosm to generate our final adjusted experimental count of CU4106 individuals. We omitted 10 total instances where adjusted counts were below 0 from our analysis, bringing the number of samples from 84 to 74. See Appendix 10 below for pertinent data.

**Appendix 10. Table of Clone Frequency Calculations**

| | temp | rep | AB +/- | proportion | CU4106 count | adjusted CU4106 count | AXS count | adjusted AXS count | total |
|---|---|---|---|---|---|---|---|---|---|
| 1 | 19 | Sample1 | NoAB | 0.514 | 5 | 5 | 0 | 0 | 5 |
| 2 | 19 | Sample2 | AB | 0.013 | 71 | 71 | 0 | 0 | 71 |
| 3 | 19 | Sample2 | NoAB | 0.514 | 20 | 18.111 | 2 | 3.889 | 22 |
| 4 | 19 | Sample3 | NoAB | 0.514 | 37 | 34.166 | 3 | 5.834 | 40 |
| 5 | 19 | Sample4 | AB | 0.013 | 53 | 53 | 0 | 0 | 53 |
| 6 | 19 | Sample4 | NoAB | 0.514 | 17 | 17 | 0 | 0 | 17 |
| 7 | 19 | Sample6 | AB | 0.013 | 49 | 49 | 0 | 0 | 49 |
| 8 | 19 | Sample6 | NoAB | 0.514 | 9 | 8.055 | 1 | 1.945 | 10 |
| 9 | 19 | Sample7 | AB | 0.013 | 76 | 76 | 0 | 0 | 76 |
| 10 | 19 | Sample7 | NoAB | 0.514 | 32 | 32 | 0 | 0 | 32 |
| 11 | 22 | Sample1 | AB | 0.586 | 73 | 72.292 | 1 | 1.708 | 74 |
| 12 | 22 | Sample1 | NoAB | 0.401 | 105 | 99.036 | 4 | 9.964 | 109 |
| 13 | 22 | Sample2 | AB | 0.586 | 77 | 74.877 | 3 | 5.123 | 80 |
| 14 | 22 | Sample2 | NoAB | 0.401 | 34 | 11.633 | 15 | 37.367 | 49 |
| 15 | 22 | Sample3 | AB | 0.586 | 12 | 9.877 | 3 | 5.123 | 15 |
| 16 | 22 | Sample3 | NoAB | 0.401 | 37 | 32.527 | 3 | 7.473 | 40 |
| 17 | 22 | Sample4 | AB | 0.586 | 22 | 2.897 | 27 | 46.103 | 49 |
| 18 | 22 | Sample6 | AB | 0.586 | 13 | 12.292 | 1 | 1.708 | 14 |
| 19 | 22 | Sample6 | NoAB | 0.401 | 13 | 11.509 | 1 | 2.491 | 14 |
| 20 | 22 | Sample7 | AB | 0.586 | 7 | 6.292 | 1 | 1.708 | 8 |
| 21 | 22 | Sample7 | NoAB | 0.401 | 21 | 21 | 0 | 0 | 21 |
| 22 | 25 | Sample1 | AB | 0.271 | 162 | 33.085 | 48 | 176.915 | 210 |
| 23 | 25 | Sample1 | NoAB | 0.559 | 254 | 129.37 | 158 | 282.63 | 412 |
| 24 | 25 | Sample2 | AB | 0.271 | 358 | 113.598 | 91 | 335.402 | 449 |
| 25 | 25 | Sample2 | NoAB | 0.559 | 179 | 73.301 | 134 | 239.699 | 313 |
| 26 | 25 | Sample3 | AB | 0.271 | 309 | 169.342 | 52 | 191.658 | 361 |
| 27 | 25 | Sample3 | NoAB | 0.559 | 192 | 104.444 | 111 | 198.556 | 303 |
| 28 | 25 | Sample4 | AB | 0.271 | 246 | 165.428 | 30 | 110.572 | 276 |
| 29 | 25 | Sample4 | NoAB | 0.559 | 261 | 212.095 | 62 | 110.905 | 323 |
| 30 | 25 | Sample5 | AB | 0.271 | 263 | 220.028 | 16 | 58.972 | 279 |

| | | | | | | | | | |
|---|---|---|---|---|---|---|---|---|---|
| 31 | 25 | Sample5 | NoAB | 0.559 | 180 | 169.746 | 13 | 23.254 | 193 |
| 32 | 25 | Sample6 | AB | 0.271 | 597 | 438.541 | 59 | 217.459 | 656 |
| 33 | 25 | Sample6 | NoAB | 0.559 | 193 | 127.53 | 83 | 148.47 | 276 |
| 34 | 25 | Sample7 | AB | 0.271 | 291 | 145.97 | 54 | 199.03 | 345 |
| 35 | 25 | Sample7 | NoAB | 0.559 | 184 | 137.461 | 59 | 105.539 | 243 |
| 36 | 30 | Sample1 | AB | 0.138 | 1445 | 902.47 | 87 | 629.53 | 1532 |
| 37 | 30 | Sample1 | NoAB | 0.448 | 263 | 235.838 | 22 | 49.162 | 285 |
| 38 | 30 | Sample2 | AB | 0.138 | 732 | 588.573 | 23 | 166.427 | 755 |
| 39 | 30 | Sample2 | NoAB | 0.448 | 244 | 202.023 | 34 | 75.977 | 278 |
| 40 | 30 | Sample3 | AB | 0.138 | 1566 | 1191.842 | 60 | 434.158 | 1626 |
| 41 | 30 | Sample3 | NoAB | 0.448 | 460 | 399.503 | 49 | 109.497 | 509 |
| 42 | 30 | Sample4 | AB | 0.138 | 848 | 93.447 | 121 | 875.553 | 969 |
| 43 | 30 | Sample4 | NoAB | 0.448 | 341 | 249.637 | 74 | 165.363 | 415 |
| 44 | 30 | Sample5 | AB | 0.138 | 328 | 128.449 | 32 | 231.551 | 360 |
| 45 | 30 | Sample5 | NoAB | 0.448 | 372 | 307.799 | 52 | 116.201 | 424 |
| 46 | 30 | Sample6 | NoAB | 0.448 | 354 | 287.33 | 54 | 120.67 | 408 |
| 47 | 30 | Sample7 | AB | 0.138 | 1908 | 1459.01 | 72 | 520.99 | 1980 |
| 48 | 30 | Sample7 | NoAB | 0.448 | 233 | 209.542 | 19 | 42.458 | 252 |
| 49 | 32 | Sample1 | AB | 0.214 | 536 | 169.46 | 100 | 466.54 | 636 |
| 50 | 32 | Sample1 | NoAB | 0.283 | 161 | 90.111 | 28 | 98.889 | 189 |
| 51 | 32 | Sample2 | AB | 0.214 | 580 | 492.03 | 24 | 111.97 | 604 |
| 52 | 32 | Sample2 | NoAB | 0.283 | 215 | 171.96 | 17 | 60.04 | 232 |
| 53 | 32 | Sample3 | AB | 0.214 | 578 | 515.688 | 17 | 79.312 | 595 |
| 54 | 32 | Sample3 | NoAB | 0.283 | 252 | 219.087 | 13 | 45.913 | 265 |
| 55 | 32 | Sample4 | AB | 0.214 | 2101 | 1873.745 | 62 | 289.255 | 2163 |
| 56 | 32 | Sample4 | NoAB | 0.283 | 121 | 77.96 | 17 | 60.04 | 138 |
| 57 | 32 | Sample5 | AB | 0.214 | 550 | 201.787 | 95 | 443.213 | 645 |
| 58 | 32 | Sample5 | NoAB | 0.283 | 109 | 60.897 | 19 | 67.103 | 128 |
| 59 | 32 | Sample6 | AB | 0.214 | 473 | 418.019 | 15 | 69.981 | 488 |
| 60 | 32 | Sample6 | NoAB | 0.283 | 259 | 183.047 | 30 | 105.953 | 289 |
| 61 | 32 | Sample7 | AB | 0.214 | 2796 | 1890.645 | 247 | 1152.355 | 3043 |
| 62 | 32 | Sample7 | NoAB | 0.283 | 201 | 157.96 | 17 | 60.04 | 218 |
| 63 | 38 | Sample2 | AB | 0.071 | 1135 | 273.863 | 66 | 927.137 | 1201 |

| | | | | | | | | | |
|---|---|---|---|---|---|---|---|---|---|
| 64 | 38 | Sample2 | NoAB | 0.076 | 795 | 380.056 | 34 | 448.944 | 829 |
| 65 | 38 | Sample3 | AB | 0.071 | 1051 | 385.576 | 51 | 716.424 | 1102 |
| 66 | 38 | Sample3 | NoAB | 0.076 | 1912 | 264.43 | 135 | 1782.57 | 2047 |

### Appendix 11. Alternative Experimental Conditions With and Without Antibiotics

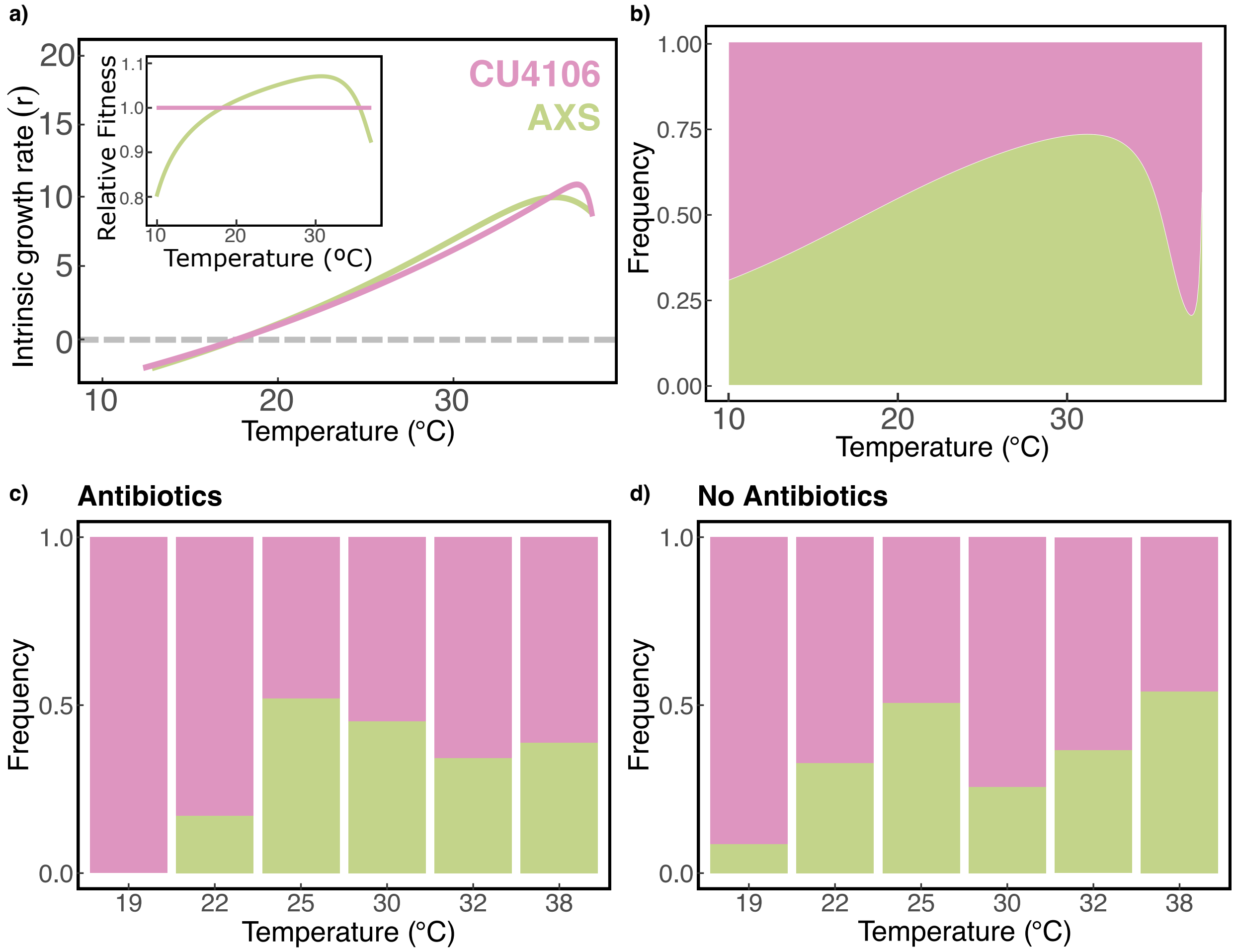

**Appendix 11 Figure S1**: a) r-TPC of strains CU4016 and AXS. Inset: relative fitness against temperature. b) Model predictions of strain frequency across temperatures. c) Observed frequencies in paromomycin-treated assays. d) Same as in c, but replicated in paromomycin-free assays.

**Appendix 12. Shape Parameter Value Estimates**

| | Strain | $CT_{min}$ | $R_{peak}$ | $E_a$ | $T_{opt}$ |
|---|---|---|---|---|---|
| 1 | A1868 | 14.12826 | 3.204581 | 0.1152172 | 33.81294 |
| 2 | B2192 III | 20.74148 | 4.047828 | 1.0803523 | 36.31143 |
| 3 | IA388 | 17.73547 | 4.417694 | 0.1508701 | 36.79342 |
| 4 | CU123 | 17.53507 | 4.689276 | 0.1557409 | 36.98405 |
| 5 | 19617-1 | 19.13828 | 4.799043 | 0.1714436 | 37.29445 |
| 6 | CU427.4 | 15.63126 | 4.809853 | 0.1437632 | 36.94217 |
| 7 | A*III | 17.13427 | 5.082249 | 0.2300526 | 34.33175 |
| 8 | CU304 | 21.34269 | 5.258839 | 0.2283879 | 36.32032 |
| 9 | A*V | 12.32465 | 5.357994 | 0.1724331 | 34.49005 |
| 10 | SB3539.1 | 15.63126 | 6.063128 | 0.2826673 | 31.36861 |
| 11 | CU438.1 | 14.92986 | 6.234928 | 0.2658260 | 34.67446 |
| 12 | B2086.2 | 17.73547 | 6.344578 | 0.2325668 | 35.66275 |
| 13 | B*VII | 17.83567 | 6.633335 | 0.2298060 | 36.64184 |
| 14 | 19877.1 | 18.43687 | 6.851586 | 0.2671804 | 35.41747 |
| 15 | CU428.2 | 16.03206 | 7.190352 | 0.2349089 | 35.27785 |
| 16 | IA280 | 17.83567 | 7.317976 | 0.2482356 | 37.19398 |
| 17 | 20441.1 | 16.33267 | 7.408803 | 0.2270542 | 36.09889 |
| 18 | SB1518 | 18.53707 | 8.363084 | 0.3068989 | 35.22339 |
| 19 | IMB6 | 21.94389 | 9.103220 | 0.3589357 | 36.52195 |
| 20 | DMCK72H | 18.63727 | 9.820957 | 0.2929556 | 37.09059 |
| 21 | CU4106 | 15.73146 | 9.866344 | 0.3077653 | 35.00748 |
| 22 | AXS | 20.14028 | 11.645950 | 0.3774932 | 36.63715 |

### Appendix 13. Shape Parameter Mean and Variance

| | Mean | SD |
|---|---|---|
| $CT_{min}$ | 17.45491 | 0.5199761 |
| $r_{peak}$ | 35.92072 | 0.2407054 |
| $T_{opt}$ | 6.720213 | 0.4852206 |
| $E_a$ | 0.2358767 | 0.01587644 |

### Appendix 14. Shape Parameter Covariances with *r*

| | Absolute value of covariance with r | l-95% CI | h-95% CI |
|---|---|---|---|
| $CT_{min}$ | 0.63 | 0.06 | 1.3 |
| $r_{peak}$ | 0.65 | 0.2 | 1.5 |
| $T_{opt}$ | 0.35 | -0.18 | 0.9 |
| $E_a$ | 0.001 | -0.15 | 0.008 |

**Appendix 15. $r_{peak}$ Stats Table**

| | Estimate | SD | P-value |
|---|---|---|---|
| Linear term | 1.81 | 0.16 | **≤0.001** |
| Quadratic term | -0.28 | 0.12 | 0.24 |
| Low Temp | -6.25 | 0.26 | **≤0.001** |
| Med Temp | -3.20 | 0.26 | **≤0.001** |
| Linear*LowTemp | -2.13 | 0.22 | **≤0.001** |
| Linear*MedTemp | -1.27 | 0.22 | **≤0.001** |
| Quadratic*LowTemp | 0.14 | 0.17 | 0.70 |
| Quadratic*MedTemp | -0.08 | 0.17 | 0.82 |

**Appendix 16. $E_a$ Stats Table**

| | Estimate | SD | P-value |
|---|---|---|---|
| Linear term | 1.61 | 0.15 | **≤0.001** |
| Quadratic term | -0.12 | 0.13 | 0.62 |
| Low Temp | -6.07 | 0.27 | **≤0.001** |
| Med Temp | -3.06 | 0.27 | **≤0.001** |
| Linear*LowTemp | -2.03 | 0.21 | **≤0.001** |
| Linear*MedTemp | -1.20 | 0.21 | **≤0.001** |
| Quadratic*LowTemp | -0.22 | 0.18 | 0.53 |
| Quadratic*MedTemp | -0.34 | 0.18 | 0.33 |

**Appendix 17. $CT_{min}$ Stats Table**

| | Estimate | SD | P-value |
|---|---|---|---|
| Linear term | 0.44 | 0.18 | **0.02** |
| Quadratic term | -0.12 | 0.13 | 0.66 |
| Low Temp | -6.15 | 0.32 | **≤0.001** |
| Med Temp | -3.21 | 0.32 | **≤0.001** |
| Linear*LowTemp | -1.19 | 0.26 | **≤0.001** |
| Linear*MedTemp | -0.88 | 0.26 | **≤0.001** |
| Quadratic*LowTemp | -0.04 | 0.19 | 0.90 |
| Quadratic*MedTemp | -0.06 | 0.19 | 0.89 |

**Appendix 18. $T_{opt}$ Stats Table**

| | Estimate | SD | P-value |
|---|---|---|---|
| Linear term | -0.32 | 0.20 | 0.11 |
| Quadratic term | -2.10 | 0.21 | **≤0.001** |
| Low Temp | -7.43 | 0.40 | **≤0.001** |
| Med Temp | -3.95 | 0.40 | **≤0.001** |
| Linear*LowTemp | 0.01 | 0.28 | 0.98 |
| Linear*MedTemp | -0.23 | 0.28 | 0.42 |
| Quadratic*LowTemp | 2.52 | 0.30 | **≤0.001** |
| Quadratic*MedTemp | 1.42 | 0.30 | **0.02** |

**Appendix 19: Adaptive Landscape with Continuous Temperature Variable**

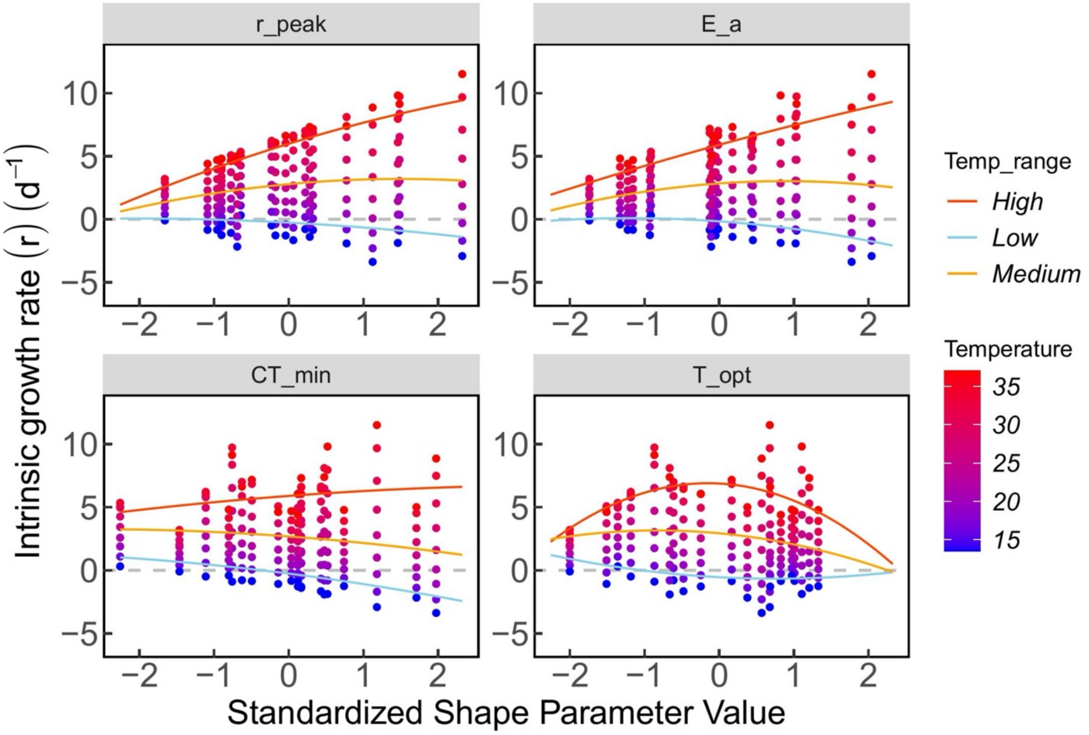

**Appendix 19 Figure S1:** Temperature across the adaptive landscape displayed as a continuous variable, provided as an alternative to the binned temperature treatments originally displayed in Figure 3.

### Appendix 20. Modeling Temperature Fluctuations

Temperature fluctuations were incorporated by sampling T at each time step from a target normal distribution that has for mean the mean of the temperature at which the model is being ran, and a set variance. Operationally, this means that the r-TPC fit was extrapolated above and below the maximum and minimum temperature for which data was collected so that temperatures could be sampled above and below 19 and 38°C and that still would result in an r estimate for that time step. However, that means that we are less confident in the effects of temperature fluctuations at colder and warmer temperatures. We assumed three variance scenarios (low, medium and high magnitude of temperature fluctuations).

Our model shows that temperature effects become increasingly strong as we allow the model to run for longer, i.e. more time steps (Fig 1a-c, d-f, g-i). While the results presented in the main text are totally robust to temperature fluctuations under medium levels of temperature fluctuations (Fig 1 d-f), they are also robust at low temperatures even at high levels of variation (Fig 1a-c). However, at high levels of temperature fluctuations we see a different pattern of response at warmer temperatures that is a direct consequence of the shape of the two TPCs considered. At very high temperatures, the r-TPC of genotype CU4016 is higher than that of AXS, but only for a limited temperature range. When temperature fluctuations are very high, and the mean temperature is contained within that range, the chance of sampling temperature outside of that range, which include temperatures where the r-TPC of CU4016 dips below of that of AXS increases, leading to a qualitative change in the evolutionary model. This means that a model like that used here is likely robust to temperature fluctuations when variation is low to moderate and will be less robust if variation is very high but also multiple r-TPC crossings are expected within a small temperature range, like that observed at high temperatures in our study system (Fig 1, Main text Fig 4d).

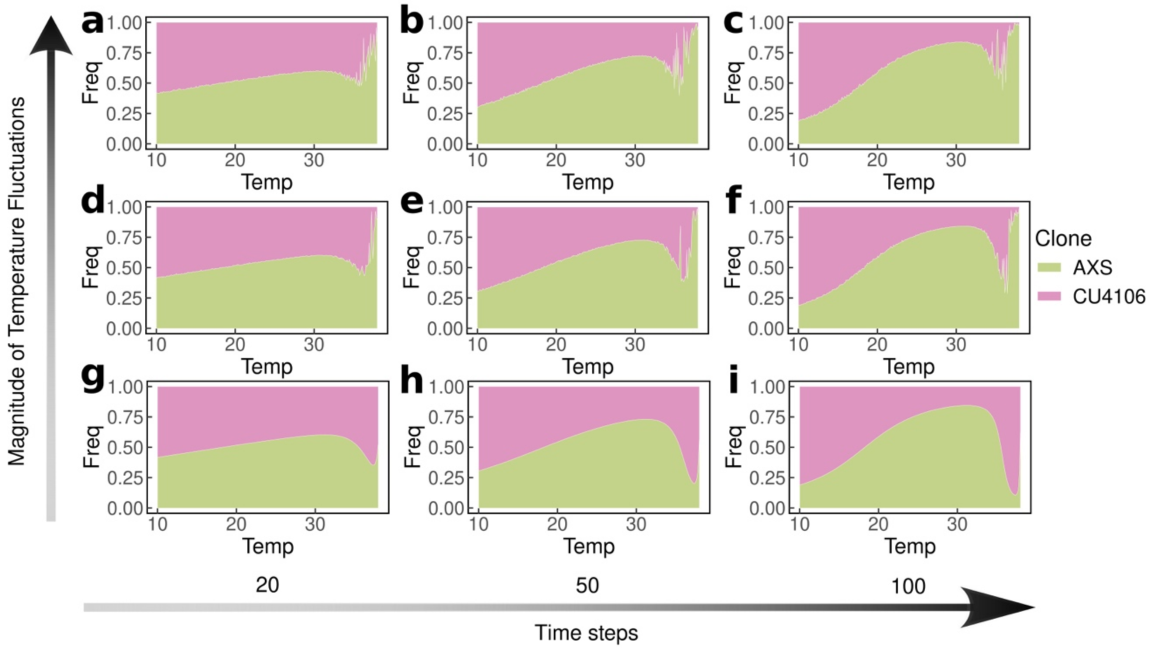

**Appendix 20 Figure S1**: Figure displaying model estimate of relative clone frequencies across temperature for 9 different scenarios (a-i) involving increasing time-steps and increasing magnitude of temperature fluctuations. Temperatures are in °C.
